# Rhythmic entrainment of saccadic eye-movements in macaque FEF

**DOI:** 10.1101/2023.05.17.540939

**Authors:** Yeganeh Shaverdi, Seyed Kamaledin Setarehdan, Stefan Treue, Moein Esghaei

## Abstract

Saccadic eye movements play a key role in gaining information about the surrounding environment. However, the neural mechanisms underlying the timing of these eye movements remain poorly understood. Here, we investigated the entrainment of saccadic eye movement by oscillatory neural activities in rhesus monkeys performing a visual foraging task. We found that saccades are phase-locked to beta LFP oscillations (16-22 Hz) in the frontal eye field (FEF), 100 ms before saccade onset, supporting a causal role of these oscillations in saccade timing. Furthermore, we show that the alignment between saccades and FEF LFPs varies, depending on the spatial relationship between the saccade target and the response field (RF) of neurons in the FEF. These findings suggest that the phase of the oscillatory neural activities determines the timing and direction of saccades.

## Introduction

Primates, including humans, use frequent saccadic eye movements (rapid shifts of gaze) to explore details of their surrounding environment [1-4]. Saccades have been shown to determine the neuronal excitability in the visual cortex by resetting the phase of neural populations [5, 6]. Saccades have also been shown to play an important role in coordinating neural communication between distributed brain areas, for instance the human amygdala and hippocampus [7].

The temporal pattern of saccades and several other active sensing modalities, such as sniffing and whisking, is rhythmic [8-13]. However, the neural processes underlying this rhythmicity and the exact timing of saccades are not well understood. We hypothesize that the rhythmic activity of neighboring neuronal population is functionally relevant for saccade alignment.

Previous studies have documented a directional interaction between saccadic eye movements and LFP rhythms in various brain areas, including the monkey primary visual cortex (V1) and hippocampus [5, 14-21]. Saccades modulate the power of LFP rhythms within alpha, beta and low-gamma bands in the monkey primary visual cortex (V1) [5, 15]. Not only the power, but the phase of rhythmic LFPs have been shown to be reset by saccades, both for human and non-human primates. These studies have demonstrated that saccades reset the phase of oscillatory activity in the visual cortex and hippocampus, particularly in the delta-theta band [14, 16, 17, 19, 20] and further electrophysiological recordings in monkeys have shown a concentration of phases following fixation onset in a wider range of frequencies, alpha (8–14 Hz), beta (14–30 Hz), and gamma (30–60 Hz) bands in the upper bank of the superior temporal sulcus [18]. While saccades are known to be critical in determining the upcoming rhythmic activity, there is limited knowledge on how rhythmic activities modulate saccades; however, see [21, 22].

In this study, we investigate the role of ongoing oscillatory neural activity in the frontal eye field (FEF), a brain area that plays a key role in guiding eye movements, in the timing of saccadic eye movements. We analyzed LFPs recorded from two macaques performing a visual foraging task to examine the potential role of the FEF’s rhythmic neural activity in saccade timing.

## Results

Two rhesus monkeys were trained to perform a foraging task (Fig. 1A), searching among five potential targets (T shape) and five distractors (+ shape) randomly arranged on the screen for the one target associated with a reward. When an animal foveated (‘fixated’) the reward-associated ‘T’ (independently selected in each trial) for at least 500 ms, it received a juice reward.

**Figure 1.**
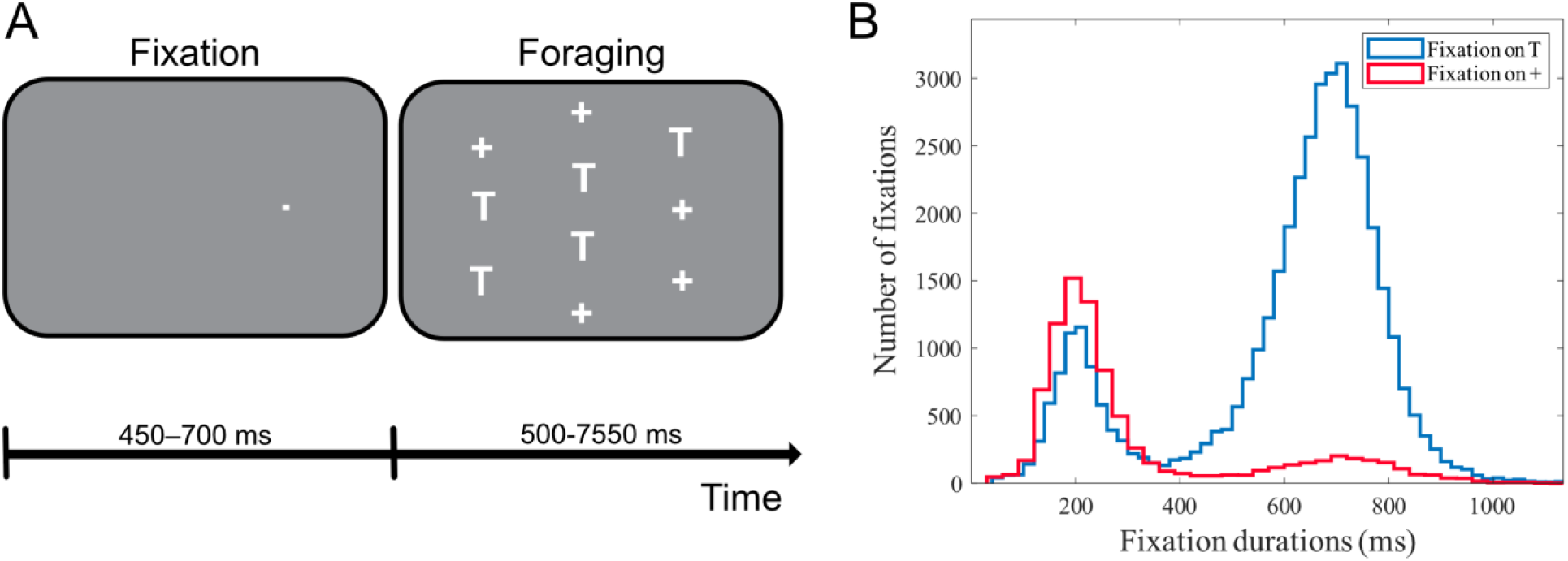
Stimulus configuration and fixation statistics. A) Example stimulus arrangement in the foraging task, with five potential targets (T) and five distracters (+). B) histogram of the duration of eye fixations on ‘T’s (blue) and ‘+’s (red), excluding the rewarding fixation.

Fixations on distractors were infrequent (17.5%, 8,933 out of 50,854) as well as substantially and significantly shorter [320 ± 221 ms (mean ± SD), p<0.0001, permutation test; n = 10,000] than fixations on potential targets (602 ± 204 ms).

LFPs and spiking activity were recorded from both animals in 61 separate sessions. There were on average 373 correctly performed trials (ending with a saccade towards the reward-associated ‘T’) in each session. In the current study, we focused on saccades which were made towards ‘T’s. The duration histogram of fixations following these saccades is plotted in Fig. 1B, showing that fixations on ‘T’s follow a bi-polar (Davies–Bouldin index [23]) distribution with two peaks at 205 and 697 ms. This suggests that the animals’ eye-fixation after making a saccade to a target lasted either a short (∼200 ms) or a long time (∼700 ms). The fact that the majority of these fixations lasted more than 500 ms (33,065 of 41,921 (78.8 %)), further indicates that the animals were correctly following the foraging paradigm.

To examine if the timing of the eye movement initiation follows an oscillatory pattern, we asked if they were coupled to the LFP signals. To this end, the average of the LFPs preceding the onset of saccades (named saccade-triggered LFP) was calculated after removal of the transient preparatory component (see Methods for details) (Fig. 2A). This saccade-triggered LFP is clearly indicative of a locking of saccades to the preceding oscillatory fluctuations of the LFP, especially within the beta band (18-21 Hz). To quantify the alignment of saccade onsets to the phase of the preceding ongoing neural oscillations, we calculated the phase-locking value (PLV) at the start of the saccades across different frequency ranges. LFPs were filtered into sweeping (step size: 1 Hz) frequency bands of 3 Hz width and a lower boundary ranging from 10 to 25 Hz. Next, we extracted the instantaneous phase of the filtered LFPs for the interval [-300, 0] ms before the saccade onset time and calculated the similarity of phases for each time-frequency pair. Fig. 2B shows the across-saccade phase similarity for all time-frequency pairs, indicating that saccades are phase-locked to the LFP oscillations for most frequencies lower than 22 Hz as early as ∼100 ms before the saccade onset. Fig. 2C plots the frequency-resolved PLV at the time of saccade onset (x-axis indicates the middle of each frequency band), showing that saccade onsets were maximally locked to the frequency band of 18-21 Hz (p<0.01, Rayleigh test; corrected for multiple comparisons using FDR). These results show that the onset of saccades is typically aligned to the phase of the oscillatory LFP, i.e. follow a rhythmic pattern.

**Figure 2.**
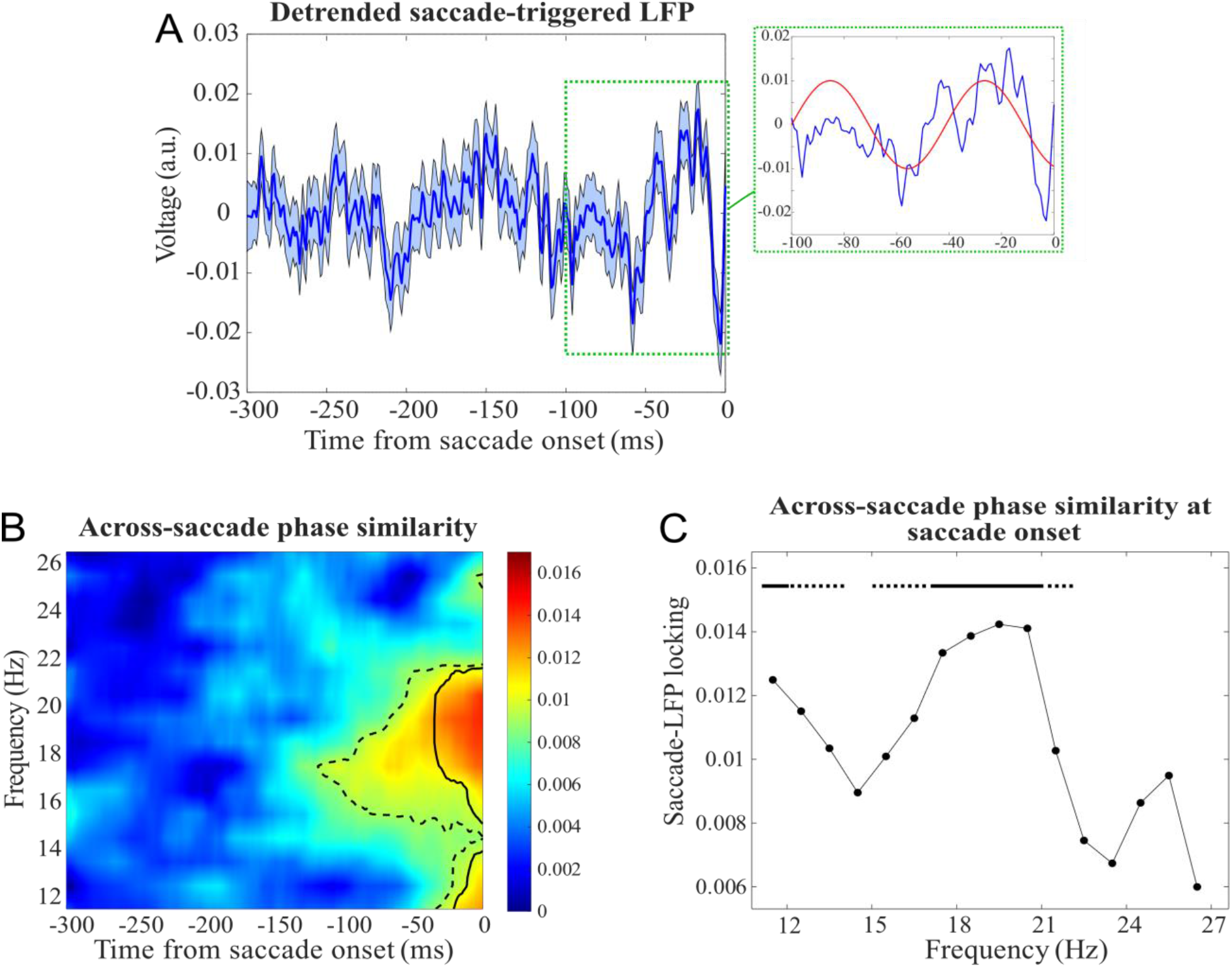
Dependence of the onset of saccades upon the phase of ongoing LFP oscillations. A) Saccade-triggered LFP across all saccades (error bars show the standard error of mean), with a zoomed-in version for [-100,0] ms showing a ∼17 Hz oscillatory component (red curve) during the interval. B) Across-saccade phase similarity; dashed and solid border lines indicate significant phase clustering values (Rayleigh test, p<0.05) before and after controlling for multiple comparisons (using FDR correction), respectively. C) Across-saccade phase similarity computed at the saccade onset for each frequency band (solid and dashed lines show significant phase clustering with a threshold of p<0.01 and p<0.05, respectively for each frequency band).

To examine the functional relevance of the locking of saccade onsets to the LFP phase, we next categorized the saccades based on their direction relative to the response field (RF); saccades towards the RF ([-45° +45°]) and saccades away from the RF ([135° 225°]) (Fig. 3). Fig. 3A and 3B show the saccade-triggered LFP for these saccades (towards and away from RF, respectively), visually clarifying how the toward-RF saccades follow a saccade-locked beta component, whereas away-RF saccades lack such coupling (see the filtered beta components overlaying the original signals). The across-saccade similarity of instantaneous LFP phases were then calculated within each group of saccades and plotted across time ([-150, 0] ms around the saccade onset) along different frequencies (Fig. 3C and 3D for saccades towards and away from RF, respectively). Saccades towards the RF exhibited a significant phase similarity starting from at least 150 ms before the saccade onset, while no such phase similarity was observed for saccades away from the RF (Fig. 3C and 3D). We also observed a gradual decrease in phase locking as saccades moved further away from the RF (see Fig. S2). To rule out the possibility that this difference was due to variations in LFP’s signal-to-noise ratio, we analyzed the pre-saccadic spectral power in ‘towards RF’ and ‘away from RF’ saccades (see Fig. S3). Interestingly, we found that the spectral power in the beta range was lower for saccades towards the RF, suggesting that differences in signal-to-noise ratio did not account for the observed difference in phase-locking (see [24] for similar results). These results suggest that not only the timing of saccades is aligned to the phase of LFPs, but also that this alignment changes as a function of the spatial relationship between the saccade target and the RF, indicating a role of oscillatory neural activities in determining the direction of saccades relative to a neuron’s RF. These findings indicate that oscillatory neural activity has a distinct role in regulating the generation of saccades and their direction relative to the RF, shedding light on the neural activity patterns underlying eye movements.

**Figure 3.**
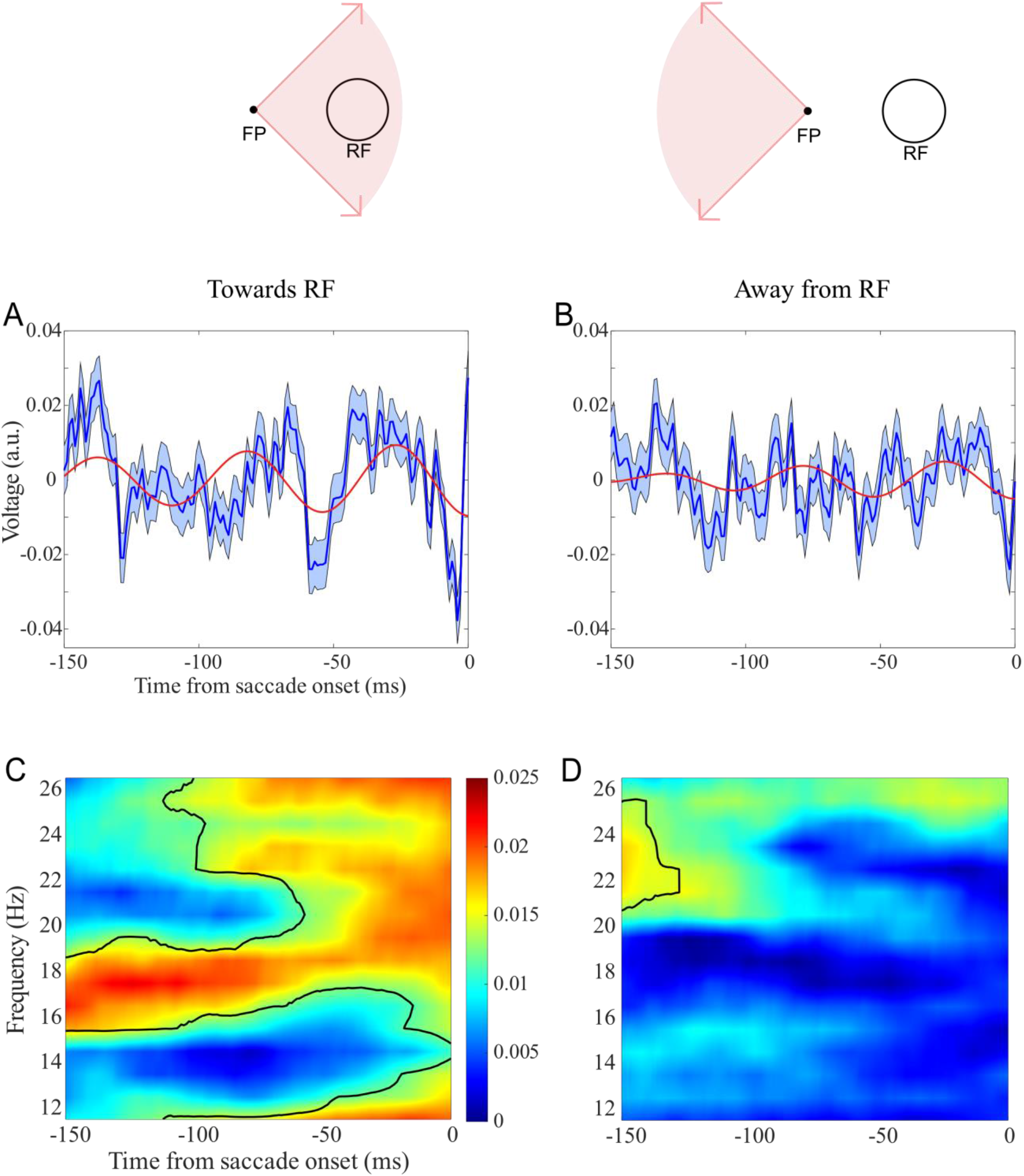
Direction dependence of across-saccade phase similarity. A) Saccade-triggered LFP for saccades towards RF and B) away from RF (error bars show the standard error of the mean). Red curves show the saccade-triggered filtered LFP (18-21 Hz band-pass filtering). C) Across-saccade phase similarity for saccades towards RF and D) away from RF (Black border lines indicate significant clusters of values (Rayleigh test, p<0.05) after controlling for multiple comparisons using FDR correction.

## Discussion

Uncovering the neural processes that govern saccadic eye movements is crucial for understanding how we selectively sample visual information from our environment. Correspondingly, there has been a growing recent interest in the relationship between eye movements and brain rhythms. Here, we show that saccades are functionally phase-locked to the beta oscillations of FEF LFPs, suggesting a specific temporal pattern in neural activity that affects the generation of saccadic eye movements. Our results shed light on the underlying neural mechanisms that govern eye movements and provide insights into the organization of the visuo-motor system.

Our data demonstrate a significant phase similarity in LFPs within the 100 ms period prior to saccades. Interestingly, this time window aligns with the time it takes (∼100 ms) to initiate and execute a saccade, even in everyday tasks like reading [22, 25]. Thus, our results not only support the notion that the brain’s processing of visual information and saccade planning are tightly coordinated, but also suggest that the phase-locking of saccades during this critical 100 ms period may play an important role in the initiation and execution of saccades.

FEF has been shown to play a crucial role in both motor planning (for eye movement) and visual attention, however it remains unclear whether the beta phase locking observed in our study is related to the motor or the attentional role of FEF. Our observation of saccades being locked to beta LFP oscillations in the FEF, confirms previous studies indicating the relevance of beta oscillations in motor areas. The beta frequency band is a well-known feature of LFPs recorded from the primary motor cortex and the supplementary motor area [26, 27], and has been shown to be crucial for motor planning, movement execution, and motor inhibition [28, 29]. This suggests that the involvement of beta oscillations in saccadic eye movements may reflect the motor planning and execution processes involved in generating eye movements. Further studies are needed to determine the potential functional significance of the beta phase locking to the guidance of pre-saccadic visual attention.

There are indications that theta rhythmic neural activities may also have a functional role in driving saccadic eye movements [30]. We speculate that beta-driven saccades may be further cross-frequency coupled to theta rhythms. This implies that beta rhythms are stronger during certain phases of theta oscillations, and weaker during other phases, potentially driving saccades relative to the theta phases. This emphasizes the fact that coupling of saccades to a certain rhythmic neural activity does not imply a rhythmicity of saccades within that frequency and raise intriguing questions about the relationship between different frequency bands and their functional roles in saccade generation. Further investigation of such cross-frequency coupling may shed light on the neural mechanisms underlying the coordination of selective visual processing and movements of the eye towards the focus of attention.

An investigation of the interaction between the ongoing oscillatory activity and saccades is only possible with a proper filtering of the oscillatory components. This becomes a particular challenge in presence of non-stationary signal components, such as those related to transient preparatory activities, causing a potential artifact distorting the conclusions drawn. In our study, we took extra care to ensure the reliability of our observations by a polynomial estimation of the transient neural activity component preceding each saccade. We are confident, that this approach has removed not only the average transient component within a neural population, but also to have captured the saccade-by-saccade variation of this component, enabling an accurate estimation of the intrinsic rather than induced oscillatory activities relevant to saccades.

Our study sheds light on the neural processes underlying saccadic eye movements. The phase-locking of saccades to beta oscillations in the FEF LFPs highlights the temporal coordination between sensory signals and the timing of eye movements. These findings contribute to our understanding of the organization of the visuo-motor system and have implications for studying the neural mechanisms involved in other sensorimotor tasks.

## Methods

We used data from the study conducted by Mirpour et al. [31] which are accessible through https://osf.io/und8y/. All experiments were approved by the Chancellor’s Animal Research Committee at the University of California–Los Angeles to comply with the guidelines established in the Public Health Service Guide for the Care and Use of Laboratory Animals. LFPs and spikes were recorded from two behaving male rhesus macaques (8–12 kg). Surgical and implantation procedures as well as localization of the region of interest (FEF) are reported in previous papers [31, 32].

### Behavioral task and electrophysiological recording

Two monkeys were trained on a visual foraging task. They started a trial by fixating on a spot that appeared on one side of the screen (Fig. 1A). After a delay of 450–700 ms, an array of five potential targets (T) and five distractors (+) was presented (each one with the size of 1.2° * 0.8°), with one over the fixation spot. One of the ‘T’s was associated with a juice reward, such that if the monkey fixated on it within 1.5° and remained fixation for 500 ms within 8 s from the start of the trial, the animal would get the reward. The stimuli were arranged so that when the monkey looked at one stimulus, the RF of an FEF neuron was likely to overlap one other stimulus. The stimuli locations, including the target, were randomly assigned among the 10 spatial locations on each trial. Neurons were excluded from the study if their RFs were so large that they would encompass two stimuli in the array. RF centers ranged from 2.8° eccentricity to 15° eccentricity, and RF sizes ranged from 1.25° to 6.5° radius in the horizontal direction and 1.25° to 4° radius in the vertical direction [31].

LFPs and single units were recorded from 61 distinct FEF sites from 2 animals while they performed the foraging task. In each session, correctly performed trials (ending with a saccade towards the reward-associated ‘T’) were selected.

### Data analysis

Scripts were programmed in MATLAB (MathWorks, Inc.). The total of 60,857 and 10,150 fixations were made onto ‘T’s and ‘+’s, respectively. We performed a permutation test to examine whether there was a significant difference between the duration distribution of the two fixation types. To this end, we selected fixation-to-’T’s before the rewarding saccade (41,921) and compared their duration to that of the fixation-to-’+’s. We selected 8,933 out of 41,921 data from the fixations landing on ‘T’s to make an equal data size of both fixation categories. Then we shuffled them with the ‘+’s fixation durations data and divided them into two equal sets with the same proportion of each fixation. We performed this shuffling for 100 iterations, calculated the mean of each category in each iteration, subtracted the means, and saved those values. The random selection of data was also made 100 times. So, the distribution of 10,000 values of the mean’s difference was created. We compared this distribution with the main difference of the distributions’ mean. The main difference was greater than 0.9999 of the distribution, suggesting that the fixations on ‘+’s were significantly (p<0.0001, permutation test; n = 10,000) and substantially shorter [320 ± 221 ms (mean ± SD)] than fixations on ‘T’s (602 ± 204 ms).

To test for the bipolarity of the fixation durations distribution (Fig. 1B), we used the Davies– Bouldin index. We created a clustering evaluation object containing data to evaluate the optimal number of data clusters. The clustering was performed using the Gaussian mixture distribution algorithm and evaluated with the Davies-Bouldin index (DBI) values as a criterion. By calculating DBI for 1 to 10 clusters, the optimal number of clusters was 2, suggesting that the histogram has a bi-polar distribution. To study the neural activity related to saccades, LFP signals within [-300,0] ms the saccade onset was analyzed. To avoid transient neural activity evoked by preceding saccades, we focused on those saccades that were at least 500 ms apart from the previous saccade. Finally, 37,027 LFP signals (within the [-300,0] ms window from the saccade onset) were selected for analysis. A notch filter was applied to the LFP signals to remove the 60 Hz power line interference. Saccade-triggered LFPs were then calculated based on frequencies above 5 Hz, filtered using a causal FFT-based 5 Hz FIR high-pass filter constructed from a Hamming window (designed in the Brainstorm MATLAB toolbox). We next removed a third-degree polynomial trend from each saccade separately. Fig. S1 shows the saccade-triggered LFP before and after detrending. Normalization of the saccade-triggered LFPs was performed saccade-wise, by calculating the z-score of the signals (mean=0 and standard deviation=1, calculated based on the interval –300 to 0 ms relative to saccade-onset).

### Extraction of the phase of LFP signals

#### Across-saccade phase similarity

To assess the coupling of saccadic eye movements to LFP oscillations, we calculated the similarity of the LFP phase at each time-frequency point preceding and relative to the saccade onset. To this end, the LFPs were filtered within different frequency bands from 10 to 25 Hz, using windows of 3 Hz width in steps of 1 Hz (i.e., 10-13 Hz, 11-14 Hz, etc.). To avoid a possible edge effect in the analysis period, we performed noise padding before the filtering, by adding 300 ms long randomly generated pink noises at the end of each LFP signal, i.e., band-pass filtering was performed on 600 ms long signals (consisting of 300 ms original LFP+300 ms long pink noise). Adding pink noise, rather than simple zero-padding ensured avoiding non-stationarities in the signal by maintaining the amplitude distribution of the signal in time. To ensure the similarity of the amplitude distribution between the noise and the original signal, we first scaled each signal to [-1,1] and then z-scored them afterwards. We generated 1,000 random noises for each LFP signal. For each repetition of noise-padding to each signal, the instantaneous LFP phase was extracted using the Hilbert transform at each time-frequency pair preceding the saccade onset. Then, the circular average across phases of the 1,000 repetitions was calculated according to the following formula:

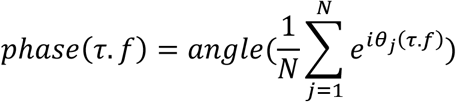

where (*τ. f*), *N*, and *θ* are a given time-frequency pair, number of repetitions, and the extracted phase, respectively. This step was performed for each of the 37,027 pre-saccadic LFP signals. Finally, the phase similarity between the obtained phases (across-saccade phase similarity) was calculated across saccades using the following equation:

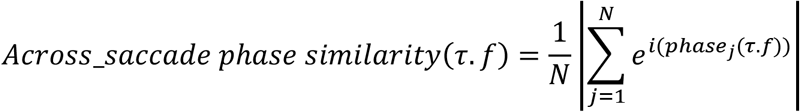

*where N* is the number of saccades. The higher the resultant value, the stronger the saccade-LFP phase coupling for a given time-frequency pair.

To check the significance of the phase similarity at each time-frequency pair (based on the value of across-saccade phase similarity), we used the Rayleigh test. The p-values were corrected for multiple comparisons using FDR correction.

#### Labeling the saccades based on their direction relative to RF

We labeled the saccades based on their angle relative to the RF of single units: saccades towards the RF ([-45° +45°]) and saccades away from the RF ([135° 225°]) (Fig. 3). To this end, for each recorded neuron we checked the direction of each saccade generated relative to the neuron’s RF. This resulted into labeling of 23,856 and 29,337 saccades to towards and away from RF, respectively. To reach a fair comparison between the two categories’ phase similarity maps, we computed the phase similarity map for the saccade away from RF case, based on a randomly selected subset of 23,856 saccades.

## Notes

https://osf.io/und8y/

### Competing Interest Statement

The authors have declared no competing interest.

