## Supplemental Figures 1, 2, 3 for "Rhythmic entrainment of saccadic eye-movements in macaque FEF"

**This file includes:**

Figs. S1 to S3

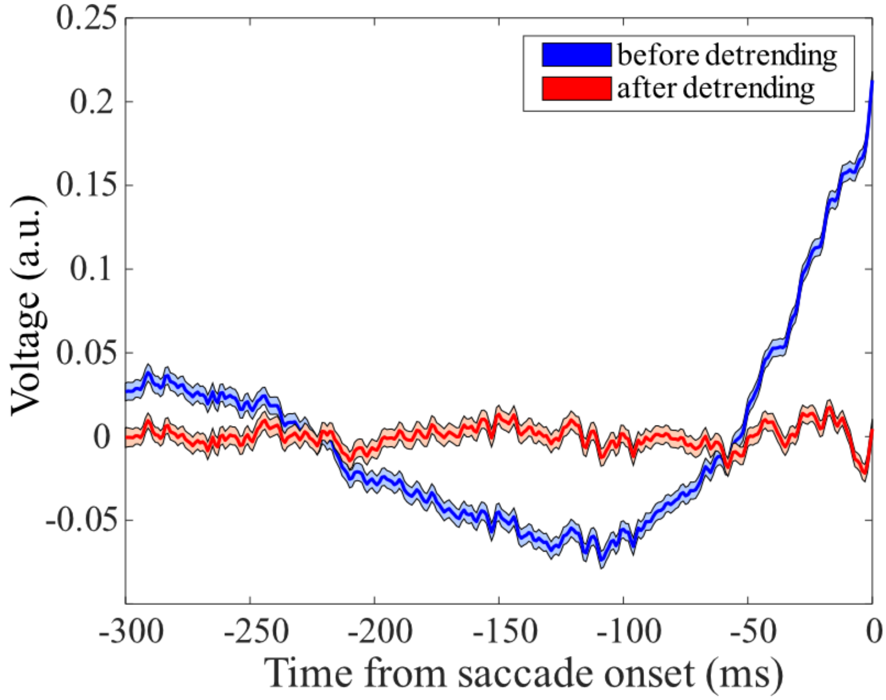


*Figure S1. Saccade-triggered LFP across all saccades before (blue) and after (red) removing the transient component before the saccade (by estimating a third-degree polynomial trend). Error bars show the SEM.*

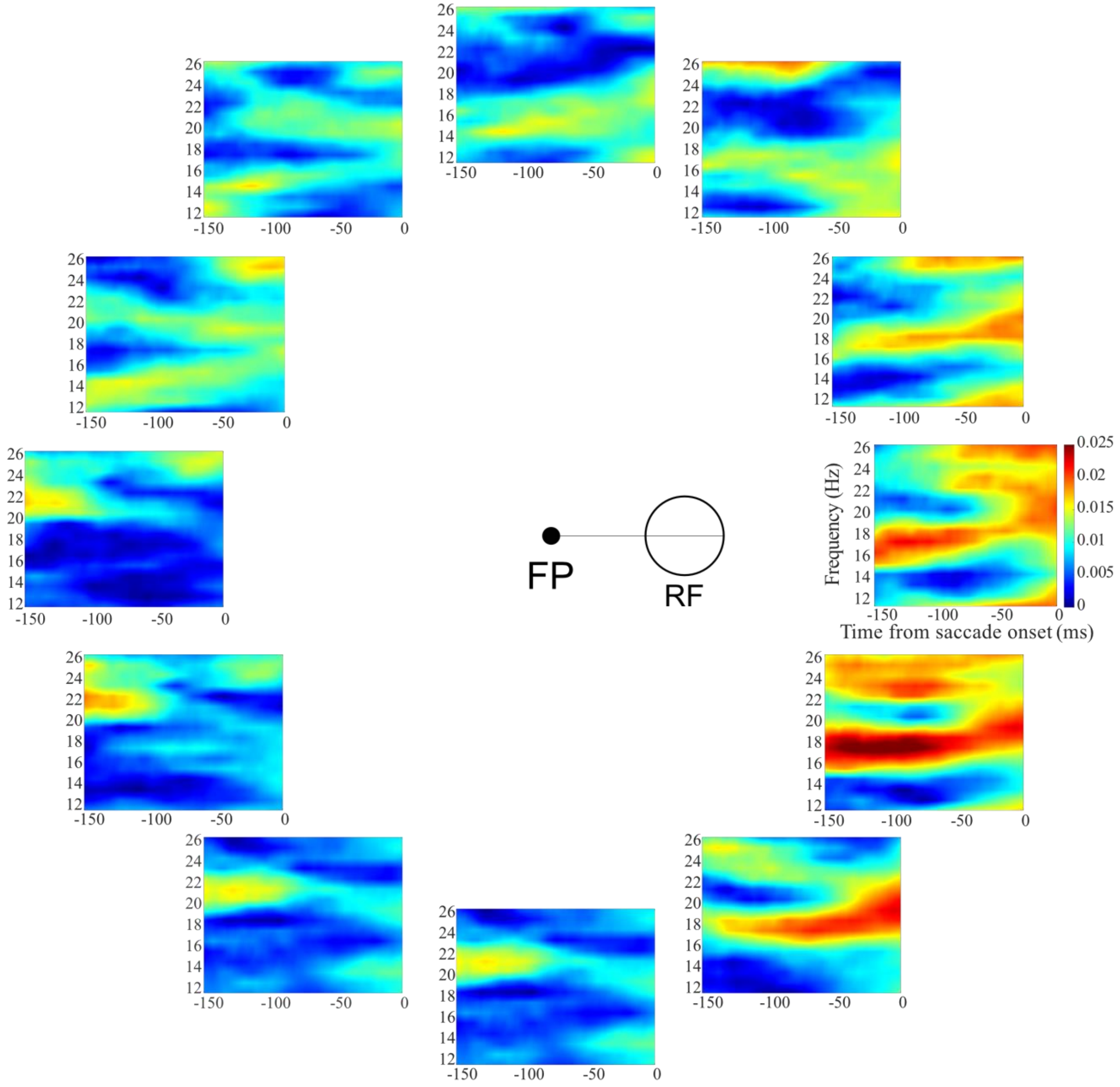


*Figure S2. Across-saccade phase similarity for various saccade directions relative to the neuron's RF. Saccades were categorized into 12 circular bins based on their direction relative to the line connecting the FP and the RF (90°-wide bins overlapping by 30°): (-45°, 45°), (-15°, 75°), ..., (-75°, -15°).*

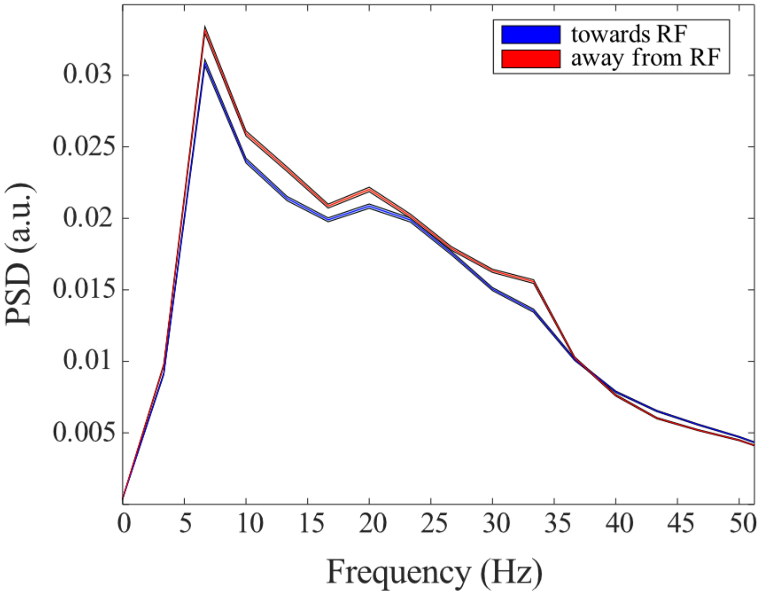


*Figure S3. Power spectral density of the pre-saccadic LFP for saccades directed towards the RF (blue) and away from the RF (red). Error bars show the SEM.*
